# Topographical relocation of adolescent sleep spindles reveals a new maturational pattern in the human brain

**DOI:** 10.1101/2021.05.25.445646

**Authors:** Ferenc Gombos, Róbert Bódizs, Adrián Pótári, Gábor Bocskai, Andrea Berencsi, Hanna Szakács, Ilona Kovács

## Abstract

Current theories of human neural development emphasize the posterior-to-anterior pattern of brain maturation. However, this scenario leaves out significant brain areas not directly involved with sensory input and behavioral control. Suggesting the relevance of cortical activity unrelated to sensory stimulation, such as sleep, we investigated adolescent transformations in the topography of sleep spindles. Sleep spindles are known to be involved in neural plasticity and in adults have a bimodal topography: slow spindles are frontally dominant, while fast spindles have a parietal/precuneal origin. The late functional segregation of the precuneus from the frontoparietal network during adolescence suggests that spindle topography might approach the adult state relatively late in development, and it may not be a result of the posterior-to-anterior maturational pattern. We analyzed the topographical distribution of spindle parameters in HD-EEG polysomnographic sleep recordings of adolescents and found that slow spindle duration maxima traveled from central to anterior brain regions, while fast spindle density, amplitude and frequency peaks traveled from central to more posterior brain regions. These results provide evidence for the gradual posteriorization of the anatomical localization of fast sleep spindles during adolescence and indicate the existence of an anterior-to posterior pattern of human brain maturation.

## Introduction

Details on the protracted maturational course of the human brain have been accumulating due to advanced brain imaging technologies^1–3^, and the characteristic back-to-front pattern has become a dominant presumption with respect to large-scale structural maturation^4^. According to the latest accounts of adolescent functional connectivity, the association areas or the so-called default mode network (DMN) of the brain are remodeled during development at an even later age than those areas that are more related to immediate sensory input and behavioral control^5,6^. While the latest maturing hot-spots in the network subserving “internal” cognition are believed to be fundamental to human evolution^7–9^, their exact connectivity and contribution to adolescent development is not known. We suggest that the investigation of spontaneous cortical activity in the absence of sensory stimulation, such as sleep, might help to move this field forward, and a more complete account of the developing large-scale functional networks will emerge.

The dynamics of large-scale brain networks are tightly linked with neural oscillatory activities. Sleep spindle oscillations are groups of 11-16 Hz waves emerging in nonrapid eye movement (NREM) sleep resulting from the rhythmic hyperpolarization rebound sequences of thalamocortical neurons^10^. Sleep spindles emerging from spontaneous neural activity during sleep reflect the underlying large-scale cortical functional networks^11,12^. A bimodal frequency and topography of sleep spindles has been described in adults. Slow (∼12 Hz) sleep spindles are frontally dominant, whereas fast (∼14 Hz) sleep spindles are of parietal/precuneal origin in adults^13,14^.

The precuneus is a major hub of the DMN, part of the complex neurocognitive network involved in the maintenance of conscious awareness, self-reflection, visuospatial integration and episodic memory^9,15^. Similar to the frontal cortex, precuneus expansion is a neurological specialization of Homo sapiens and is assumed to be associated with recent cognitive specializations^16^. The precuneus has been characterized by increasing functional segregation from the frontoparietal network between 8 and 26 years of age^17^. In addition to the late maturation of frontal lobe functions characterized by the well-known back-to-front maturational pattern of the brain^2–4,18^, the precuneus is also a subject of significant developmental changes in adolescence.

Based on the relatively late functional segregation of the precuneus from the frontoparietal network, as well as on the bimodal prefrontal versus precuneal sources of slow and fast sleep spindles in the adult brain, we postulate that sleep spindles undergo a topographical transformation characterized by increasing anatomical segregation during adolescent development. We assume that slow and fast sleep spindles are gradually relocated from the more central toward the prefrontal and parietal regions, respectively. Age-related topographical transfer is assumed to reflect the increasing differentiation and segregation of the frontal and precuneal subsystems in the DMN and the associated late development of phylogenetically new neuroanatomical structures in humans.

Here, we focus on the late maturation of slow frontal and fast parietal sleep spindle features in adolescence by using HD-EEG recordings in three subgroups of different ages (12, 16 and 20 years). We hypothesize that the two types of sleep spindles index distinct neurodevelopmental processes during the course of adolescent brain maturation and functional specialization. We believe that focusing on the bimodal antero-posterior source and frequency distribution of sleep spindle oscillations^13,14^, contrasting the unimodal, frontal maxima of slow waves^19,20^ provides us with a better understanding of adolescent remodeling of the human brain.

## Results

We recorded 128 channel full-night sleep HD-EEG data and detected individual-specific slow and fast spindles by the Individual Adjustment Method (IAM, see Methods/Procedure and ref.^21^). We analyzed the derivation-specific topographical distribution of the spindle parameters (density, duration, maximum amplitude and peak frequency) for NREM sleep. The distribution of the spindle parameters along the midline derivations (from Fpz to Iz) was calculated, and the maximum position of the distribution for every spindle parameter was determined. We scrutinized anterior-posterior shifts of the maximum locations as a function of age.

Here, we first describe the topographical distribution and age-related topographical changes of slow and fast spindle parameters (Figures 1 and 2). Second, we present evidence for age-related antero-posterior shifts in sleep spindle parameters (Figure 3).

**Figure 1.**
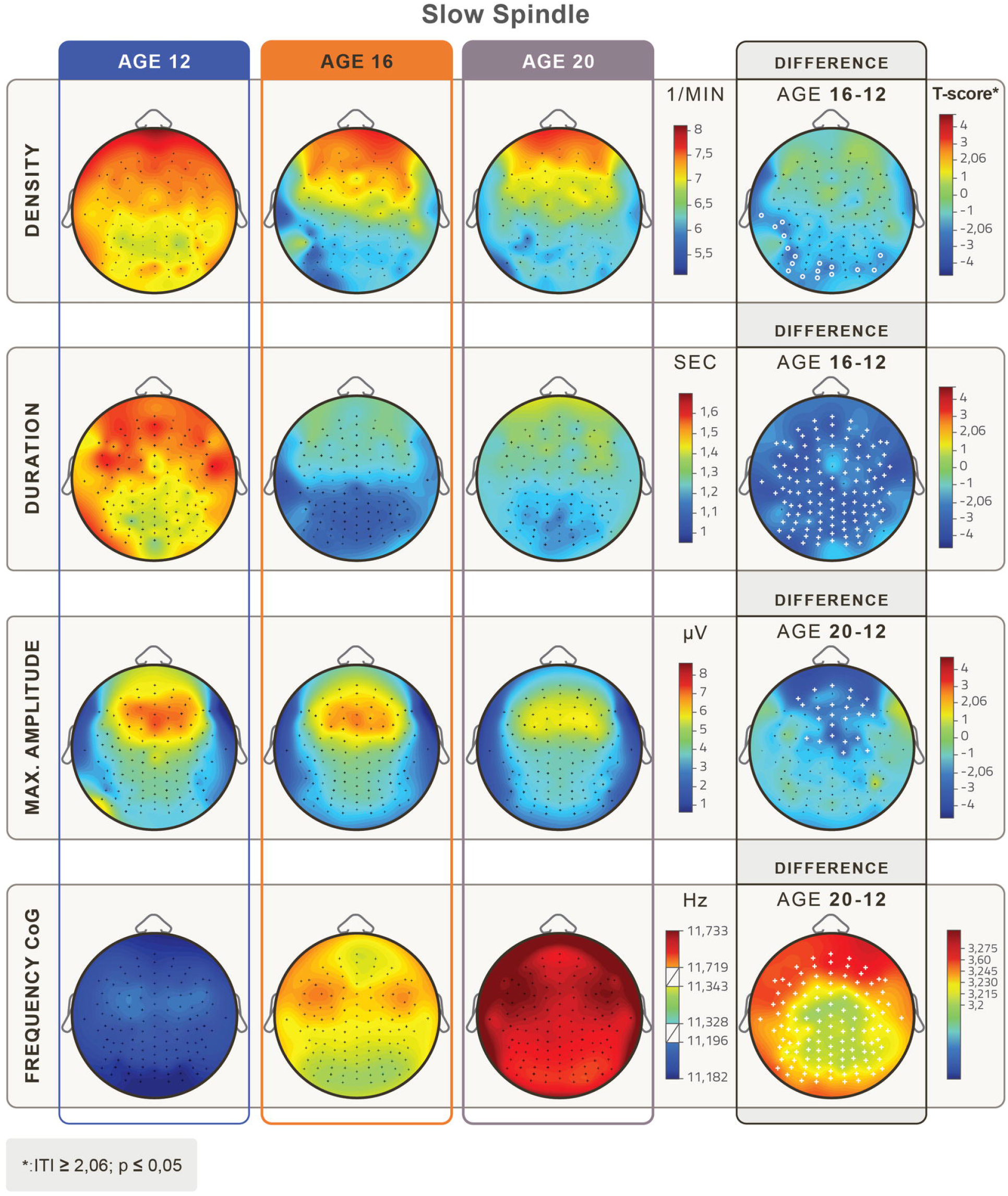
Topographical distribution of mean slow sleep spindle parameters in the age groups of 12, 16 and 20 years of age. Rows depict age-related changes in slow sleep spindle density (1/min), duration (s), maximum amplitude (μV), and frequency based on the center of gravity (CoG) (Hz). Relevant age-group differences are depicted in the 4th column. White crosses depict Rüger area significant differences and white circles depict significant but not Rüger area significant differences. Age-related decreases in density are significant with an occipital and left temporal focus. There was a significant overall decrease in duration and a frontal-frontopolar decrease in maximum amplitude. Frequency increases at all derivations with a frontal focus from the younger to the older age groups. (Level of significance: p <.05, T-score values are matched with the colorbar, T ≥ |2.06| p<0.05, at T ≥ |2.787| p<0.01 and at T ≥ |3.725| p<0.001.)

**Figure 2.**
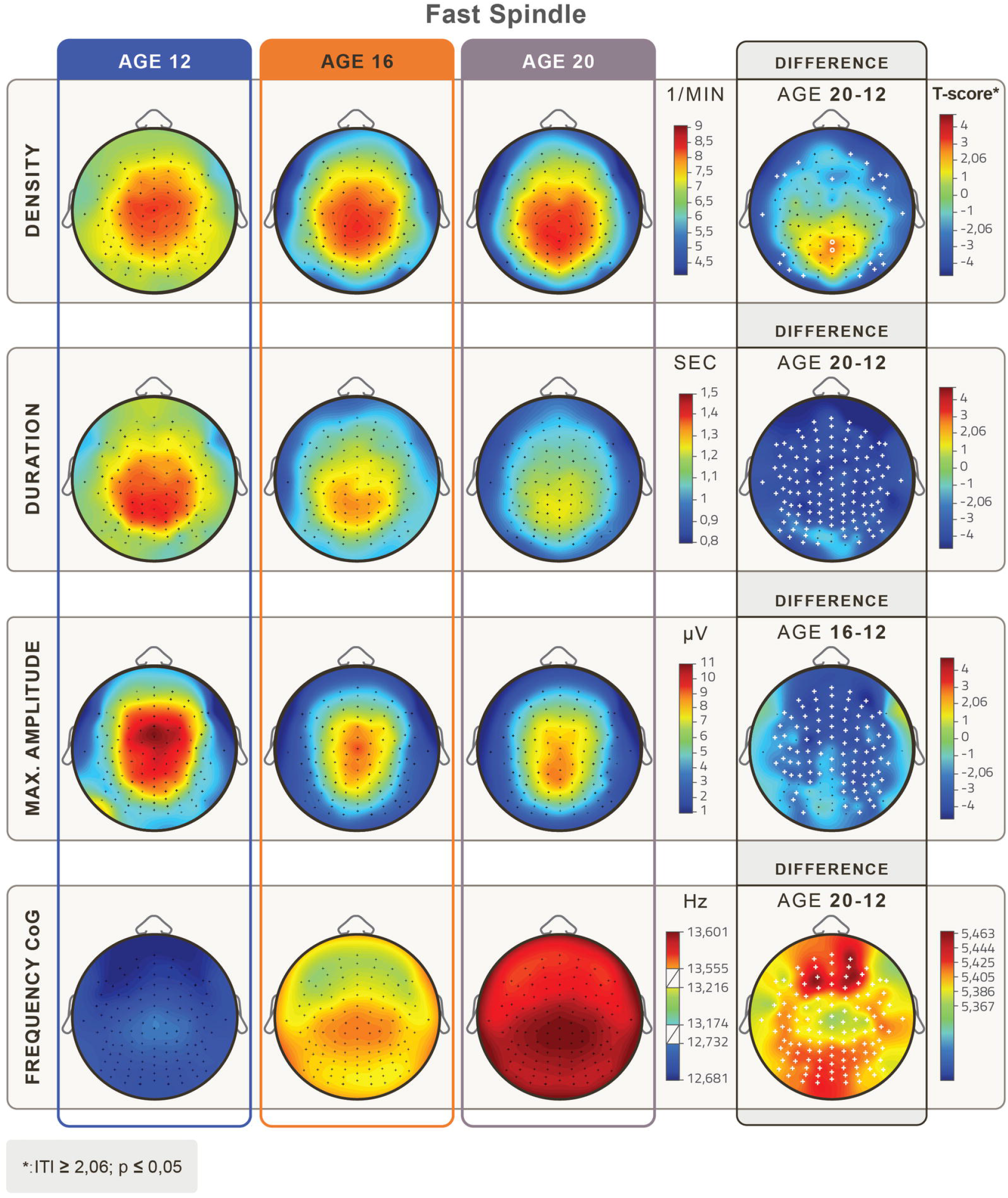
Topographical distribution of mean fast sleep spindle parameters in the age groups of 12, 16 and 20 years of age. Rows depict age-related changes in fast sleep spindle density (1/min), duration (s), maximum amplitude (μV), and frequency based on the center of gravity (CoG) (Hz). Relevant age-group differences are depicted in the 4th column. White crosses depict Rüger area significant differences, and white circles depict significant but not Rüger area significant differences. Fast spindle density increased significantly, but not Rüger significantly centro-parietally between 12 and 20 years of age, and Rüger significantly decreased at the perimeter of the scalp. There was a significant overall decrease in duration and a frontal decrease in maximum amplitude. Frequency significantly increased at all derivations with frontocentral maxima from the younger to the older age group. (Level of significance: p <.05, T-score values are matched with the colorbar, T ≥ |2.06| p<0.05, at T ≥ |2.787| p<0.01 and at T ≥ |3.725| p<0.001.)

**Figure 3.**
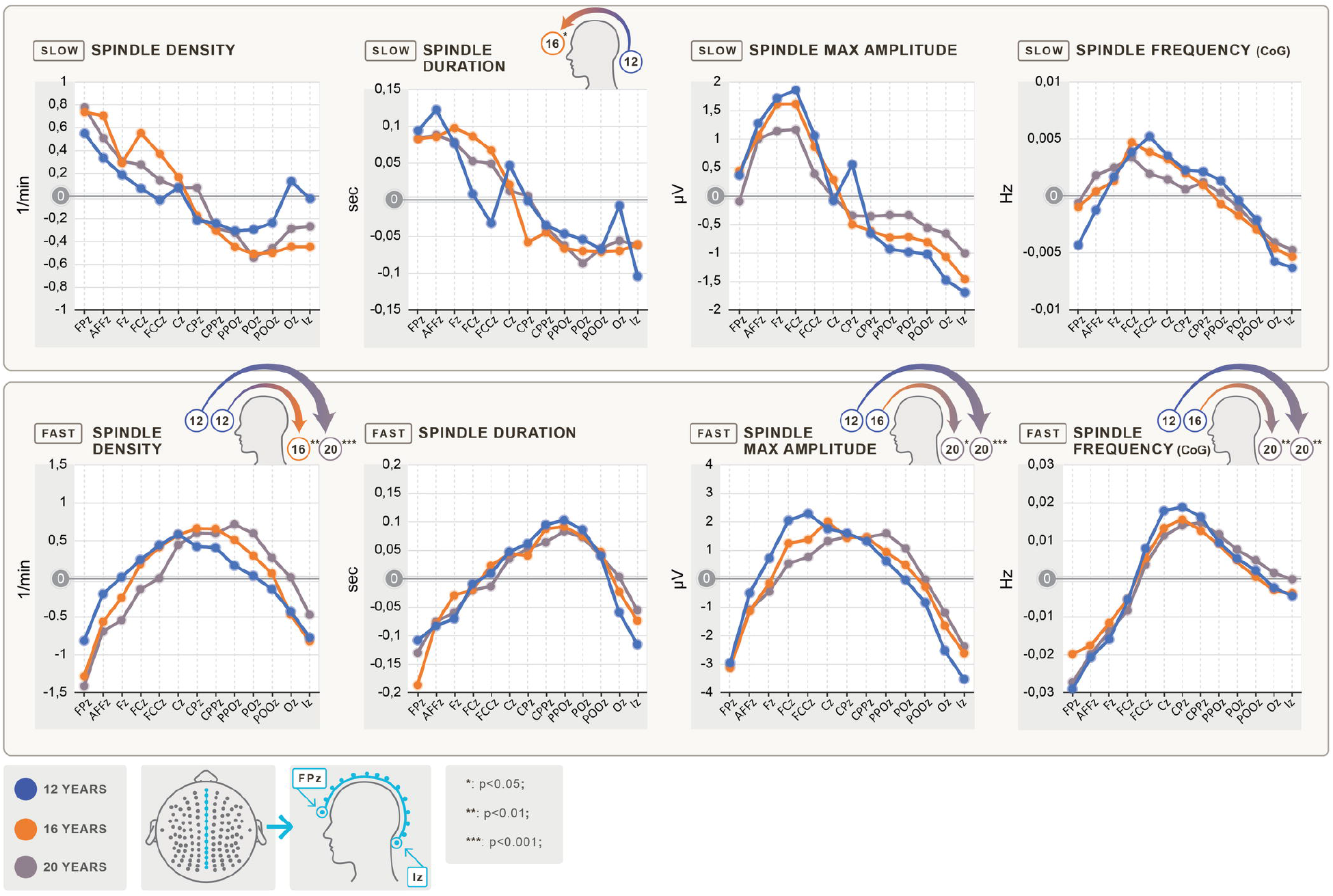
Age-related antero-posterior shifts in sleep spindle parameters. The spindle parameter values (y-axis) for the midline electrodes (Fpz to Iz on the x-axis, from left to right from frontal to occipital) are plotted by age groups. As the differences are significant but small (differences due to spatial position are much smaller than differences between age groups), the y-values display the difference from the mean value of the midline electrodes of the specific age group. The upper row depicts slow spindle parameters, while the lower row depicts fast spindle parameters. The different columns show different spindle parameters, such as density (1/min), duration (sec), maximum amplitude (μV) and frequency (Hz), based on the center of gravity (CoG). The small head figures above the graphs indicate significant differences and the direction of the difference in the positions of the maximum values between the specific age groups. Blue indicates 12-, orange indicates 16- and gray indicates 20-year-old subjects. Slow spindle parameters tended to shift toward the anterior region, whereas fast sleep spindle parameters tended to shift toward the posterior regions with age. The shift of the maximum values of slow spindle duration toward frontal regions was statistically significant. In terms of fast spindle density, maximum amplitude and frequency, the shift of the maximum values from central to centroparietal regions was statistically significant.

### Topographical distribution and age-related topographical changes of slow and fast spindle parameters

The age-dependent topographical distributions of slow and fast spindle parameters are shown in Figures 1 and 2, respectively.

In terms of duration, the decrease in slow spindles from age 12 to 16 was Rüger area significant for all recording locations (T=(−2.05 – -3.29), p= (.002 – .04)). Maximum amplitude values of slow spindles decreased with frontal and frontopolar foci between all age bins from 12 to 20 years (T=(−2.03 – -2.99), (p<.001 – .048)). Frequency increased Rüger area significantly at all derivations with a frontal focus throughout the entire adolescent period (T=(3.2 – 3.28), p= (.002 – .003)), see Figure 1.

Fast spindle density decreased Rüger area significantly at the perimeter of the scalp (T=(−2.07 – -3.06), p=(.04 – .004)). Fast spindles revealed a significant decrease in duration for all recording derivations over the entire adolescent period from age 12 to 20 (T=(−2.06 – -4.54), (p<.001 – .047)). Fast spindle maximum amplitude values decreased significantly in all areas with a frontal and central focus (T=(−2.05 – -4.54), (p<.001 – .048)). Fast spindle frequency values monotonously increased with age. The differences are Rüger area significant for the entire cortex in all age bins (T=(5.37 – 5.46), p< .001), see Figure 2.

### Age-related shifts in sleep spindle parameters

Topographical shifts of slow and fast spindle parameters can be visually assessed in the topographical distribution maps of Figures 1 and 2, respectively. While slow spindle parameters tended to shift toward anterior regions, fast sleep spindle measures seemed to shift toward posterior regions with age. To more precisely quantify these observable tendencies, we calculated the distribution of spindle parameters along the midline derivations (from Fpz to Iz) and determined the maximum position of the distribution for every spindle parameter.

Maximum values of slow spindle duration shifted toward frontal regions. The difference in the positions between the age groups of 12 and 16 years of age was statistically significant (F(1,38)=4.3; p=.045; ANOVA). In terms of fast spindle density, the difference in positions of maximum values was significant in the age groups of 12 to 16 (F(1,38)=11.27; p=.002; ANOVA) and 12 to 20 (F(1,38)=23.84; p<.001; ANOVA), revealing a central to centroparietal shift with age. Maximum values of the positions of fast spindle maximum amplitudes shifted significantly from central to centroparietal regions in the age groups of 16 to 20 (F(1,38)=4.5; p=.04; ANOVA) and 12 to 20 (F(1,38)=13.2; p<.001; ANOVA) but not between ages 12 to 16. Maximum values of the positions of fast spindle frequency also shifted significantly from central to centroparietal regions in the age groups of 16 to 20 (F(1,38)=11.76; p=.001; ANOVA) and 12 to 20 (F(1,38)=9.77; p=.003; ANOVA) (Figure 3.).

## Discussion

Here, we report evidence for distinct shifts in the predominant location of slow and fast sleep spindle oscillations during adolescent development. We found evidence for slow sleep spindle duration maxima traveling from central to anterior brain regions and fast spindle density, amplitude and frequency peaks traveling from central to more posterior brain regions. These findings coincide with reports of an age-related increase in the functional segregation of the precuneus (a major source of fast sleep spindle oscillations in humans) from the frontoparietal region from childhood to adulthood^17^.

In the late maturing frontal and parietal cortical areas, the short-range local connections are dominated by long-range functional networks during development^22^. In the course of adolescence, the anterior-posterior connectivity of the frontal and parietal lobes increases in the DMN^23,24^. Although functional connectivity may be reduced in the DMN in deep sleep stages^25,26^, it is preserved to such an extent that oscillatory activity-related functional connectivity between anterior and posterior areas corresponds to DMN characteristics during sleep spindling^27^.

Previous HD-EEG reports, based on sleep slow waves, have shown the anteriorization of sleep state-dependent neural oscillations during human ontogenesis (including the period of adolescence)^18^; however, age-related posteriorization of any frequency band has not been shown before. Thus, the late maturation of the frontal lobes is clearly supported by findings of former sleep neurophysiological studies, as well as our present report on slow sleep spindle oscillation; however, the phylogenetically new precuneal region has not yet been targeted by such studies. Although it could have been assumed that the late maturing precuneus is characterized by age-related posteriorization of sleep-state-specific oscillations, indexing the timing of developmental plasticity, such findings have been obscured by the insufficient frequency resolution of former reports.

Studies investigating the development of sleep spindles in children or adolescents do not cover the age-dependent topographical maturation of sleep spindles due to the low number of EEG electrodes^28–30^. Here, we use an individually adjusted frequency analysis of sleep spindle oscillations, with high frequency resolution and 128 channel HD-EEG recording, which is sufficiently sensitive to capture this process. With these more sensitive methods, we have been able to show that individual-specific fast sleep spindle features unequivocally travel to more posterior areas during adolescence. Considering the precuneal origin of fast sleep spindles in adult human subjects^13,14^, it seems that individualized sleep spindle analysis is a forceful tool to detect the dominant anatomical directions of brain maturation in humans.

Prevailing concepts of complex neural development and brain maturation emphasize the posterior-to-anterior or back-to-front pattern^2–4^ as an unequivocal ontogenetic feature, finding its roots in the late maturing frontal lobes. The back-to-front pattern has received substantial support from HD-EEG studies examining the age-related changes in the topography of sleep slow waves^18^. However, the above scenario of unidirectional developmental and anatomical route disregards the major hub of the DMN, which is a human-specific and late maturing neural structure of the brain, undergoing significant differentiation during adolescence. In addition to slow waves, sleep spindles are the major hallmarks of NREM sleep, known to be involved in neural plasticity with major sources in the precuneus, at least in healthy adult human subjects. Here, we provide evidence for a backward move in the anatomical localization of fast-type sleep spindles; that is, the predominantly central maxima in density and amplitude at the age of 12 years gradually become replaced with parietal regions by the age of 20 years. Similar changes, albeit at a smaller anatomical distance, take place in terms of fast sleep spindle frequency. These findings complete the back-to-front pattern of brain maturation and cohere with reports of increased differentiation of the precuneus during adolescence. To the best of our knowledge, this is the first report of an age-related front-to-back transfer of the maxima of a neural plasticity-related sleep EEG index.

Apparently, adolescence is a timeframe of significant changes in the EEG spectra, including shifts in sleep spindle characteristics^28–33^. Sleep spindles have outstanding relevance from a clinical point of view since adolescence-related psychiatric disorders such as schizophrenia^34^, ADHD^35^ and depression^36^ have all been associated with anomalies in sleep spindles. It has also been suggested that sleep spindle abnormalities are an endophenotype of schizophrenia^34,37^. Moreover, these disorders can also be characterized by cognitive impairments^38–40^. It has been demonstrated that sleep spindles and cognitive function are strongly associated^41,42^, and as such, sleep spindle abnormalities might provide a key to interpreting cognitive malfunction^10^. Further investigating the mechanisms and developmental trajectory of sleep spindles in adolescence could lead to a better understanding and novel treatment of the aforementioned disorders.

In conclusion, we focused on the late maturation of individually adjusted slow-frontal and fast-parietal sleep spindle features in adolescence by using HD-EEG recordings. Our hypothesis assuming that the two sleep spindle types might index distinct neurodevelopmental processes during the course of adolescent brain maturation and functional specialization has been confirmed by the data. We are convinced that focusing on the bimodal antero-posterior source and frequency distribution of sleep spindle oscillations has taken us closer to a better understanding of adolescent remodeling of the human brain.

## Methods

### Participants

We recruited 60 adolescent and young adult participants (30 females and 30 males, mean age 16.55 ± SD 3.70) via social media in three age groups, with an equal number of females and males in each age group: 12-year-olds (n=20, mean age=12.45 ± SD .57 years), 16-year-olds (n=20, mean age=15.91 ± SD .48 years), and young adults referred to as 20-year-olds (n=20, mean age=21.29 ± SD .51 years). Subjects with an existing neurological condition or sleep disorder were not included in the study. All subjects received a voucher in the value of HUF 20,000 (cca. USD 70.00) for their participation. The study was approved by the Ethical Committee of the Pázmány Péter Catholic University for Psychological Experiments. Adult participants and parents or legal guardians of the underage participants signed informed consent for the participation in the study according to the Declaration of Helsinki.

### Procedure

We performed 128 channel HD-EEG polysomnographic (PSG) sleep recordings of two consecutive nights in the Sleep Laboratory of Pázmány Péter Catholic University Budapest. We requested participants to maintain a regular sleep-wake schedule for 5 nights before recordings, but we did not monitor compliance. We asked participants not to consume any drugs other than contraceptives and coffee and to refrain from napping on the afternoons of the days of the experiment. Participants with any history of sleep problems or mental disorder or any neurological or medical condition were excluded. We verified such criteria by personal interviews and questionnaires completed by the parents or young adult participants themselves. We applied electrodes for electroencephalography (EEG), electromyography (EMG) and electrooculography (EOG). After checking the electrodes for skin contact and impedance levels, all participants went to sleep in the laboratory between 10.00 p.m. and 11.30 p.m. according to their preference. They slept until they awoke spontaneously.

We used a built-in EEG channel-Quick Cap (Compumedics, Australia) with 128 electrodes in three different head sizes to record EEG signals. We roughly evenly spaced the passive Ag/AgCl electrodes down to line T9-Iz-T10. The monopolar channels were referenced to a frontocentrally placed ground electrode. We applied bipolar EOG and EMG channels to measure eye movements and muscle tone, respectively, and placed the EOG channels below and to the left of the left eye (1 cm below the left outer canthus) and above and to the right of the right eye (1 cm above the right outer canthus). We determined the locations of the EMG electrodes in accordance with the recommendations of the American Academy of Sleep Medicine (AASM)^43^. We recorded the data using a BQ KIT SD LTM 128 EXPRESS (2 × 64 channels in master and slave mode, Micromed, Mogliano Veneto, Italy) recording device and subsequently visualized, saved and exported them using System Plus Evolution software (Micromed, Mogliano Veneto, Italy). We recorded all HD-EEG, EOG and EMG data with an effective sampling rate of synchronous 4096 Hz/channel with 22 bit equivalent resolution and prefiltered and amplified them with 40 dB/decade high- and low-pass input filters of 0.15–250 Hz. We applied a digital low pass filter at 231.7 Hz and downsampled the data to 512 Hz/channel by firmware.

From the recorded nighttime EEG recordings, the current analysis deals with the second night data as that was free from adaptation effects (first night effect). We manually scored sleep states of 20 sec epochs of whole night NREM EEG recordings by visual inspection according to standardized criteria^44^ and revised hypnograms. Then, we removed artifacts on four-second long epochs and scanned the whole-night recordings four times to overcome the limitations of the capacity of the visual inspector (only 32 channels of the 128 were visually inspected in a run), and finally, we excluded any epoch marked as artifact in any run from further analysis.

We detected individual-specific slow and fast spindles by using the Individual Adjustment Method (IAM)^21^. This method is based on the average amplitude spectra of NREM sleep. The frequency criteria of slow and fast sleep spindles were derived from the individual-specific peaks of the spectra between 9 and 16 Hz based on the inflexion points. We tested the slow and fast sleep spindle frequencies for frontal and centroparietal dominance, respectively. We determined the amplitude criteria for slow and fast spindles in an individual-and derivation-specific manner by multiplying the number of intraspindle frequency bins by the mean amplitude spectrum values corresponding to lower and upper frequency limits. Then, we applied a bandpass filter on the EEG for the individual slow and fast sleep spindle frequencies using a fast Fourier transformation filtering method and calculated the precise envelopes of the filtered signals. EEG segments corresponding to the envelopes transcending the amplitude criteria for at least 0.5 s are considered spindles. A scheme of spindle detection is provided in Fig. 1 of Ujma et al., 2015^45^. In fact, these segments contribute to the individual- and derivation-specific lower- and higher-frequency spectral peaks between 9 and 16 Hz. Based on the IAM approach, we determined individual and derivation-specific densities (spindles × min−1), durations (s), and amplitudes (μV) of slow, frontally dominant and fast, centro-parietally dominant sleep spindles.

We determined the peak frequency using the center of gravity (CoG) of the individualized spectral peak frequencies in the spindle range (11-16 Hz). We obtained values of CoG by using the method described in the study conducted by Dustman^46^ We excluded outliers from the original data set by using Tukey’s fences method^47^.

We analyzed the derivation-specific topographical distribution of spindle parameters and their age-dependent differences. We used a t-test to compare the activity of the corresponding electrodes between each group and performed multiple comparison correction using a modified version of the Rüger area method. We calculated Rüger area global significance as follows: 1/3 of the descriptive significance at p=.05/3 =.017 AND/OR half of descriptive significance at p=.05/2=.025. We used both criteria simultaneously (the “AND” operator) in this study^48,49^.

Finally, we calculated the distribution of the spindle parameters along the midline derivations (from Fpz to Iz) and determined the maximum position of the distribution for every spindle parameter. We examined the anterior-posterior shifts of the maximum locations as a function of age. Statistical comparisons were conducted by one-way ANOVA.

For the processing of sleep data (sleep state classification, artifact rejection and spindle detection with IAM), we used the FerciosEEGPlus 1.3 program^45^. We used Microsoft Excel for the calculation of CoG values and Rüger area corrections. We conducted all statistical tests using TIBCO STATISTICA 13.5.0.17 software (TIBCO Software Inc., 2018).

## Data availability

The datasets analyzed during the current study are available from the corresponding author on reasonable request.

## Acknowledgments

This research was supported by the Hungarian National Research, Development and Innovation Office grants NK-104481 and K-134370 to I.K. and K-128117 to B.R. and by the Higher Education Institutional Excellence Program of the Ministry of Human Capacities in Hungary within the framework of the neurology thematic program of Semmelweis University. The authors thank T. Jáger for generating the figures. We also thank the adolescents and young adults who participated in this project and Zoltán György who assisted at all sleep recordings.

## Author contributions

IK, FG and RB designed the experiments; FG, AP and GB analyzed the data; all authors contributed to writing the paper.

## Additional information Competing interests

The authors declare no competing interests.

## References

1. Paus, T. et al. Maturation of white matter in the human brain: a review of magnetic resonance studies. Brain Res. Bull. 54, 255–266 (2001).

2. Giedd, J. N. Structural Magnetic Resonance Imaging of the Adolescent Brain. Ann. N. Y. Acad. Sci. 1021, 77–85 (2004).

3. Sowell, E. R. et al. Mapping cortical change across the human life span. Nat. Neurosci. 6, 309–315 (2003).

4. Lenroot, R. K. & Giedd, J. N. Brain development in children and adolescents: insights from anatomical magnetic resonance imaging. Neurosci. Biobehav. Rev. 30, 718–729 (2006).

5. Váša, F. et al. Conservative and disruptive modes of adolescent change in human brain functional connectivity. Proc. Natl. Acad. Sci. 117, 3248–3253 (2020).

6. Whitaker, K. J. et al. Adolescence is associated with genomically patterned consolidation of the hubs of the human brain connectome. Proc. Natl. Acad. Sci. U. S. A. 113, 9105–9110 (2016).

7. Buckner, R. L. & Krienen, F. M. The evolution of distributed association networks in the human brain. Trends Cogn. Sci. 17, 648–665 (2013).

8. Uddin, L. Q., Yeo, B. T. T. & Spreng, R. N. Towards a Universal Taxonomy of Macro-scale Functional Human Brain Networks. Brain Topogr. 32, 926–942 (2019).

9. Cavanna, A. E. & Trimble, M. R. The precuneus: a review of its functional anatomy and behavioural correlates. Brain 129, 564–583 (2006).

10. Fernandez, L. M. J. & Lüthi, A. Sleep Spindles: Mechanisms and Functions. Physiol. Rev. 100, 805–868 (2020).

11. Spoormaker, V. I., Czisch, M., Maquet, P. & Jäncke, L. Large-scale functional brain networks in human non-rapid eye movement sleep: insights from combined electroencephalographic/functional magnetic resonance imaging studies. Philos. Trans. R. Soc. Math. Phys. Eng. Sci. 369, 3708–3729 (2011).

12. McVea, D. A., Murphy, T. H. & Mohajerani, M. H. Large Scale Cortical Functional Networks Associated with Slow-Wave and Spindle-Burst-Related Spontaneous Activity. Front. Neural Circuits 10, (2016).

13. Anderer, P. et al. Low-resolution brain electromagnetic tomography revealed simultaneously active frontal and parietal sleep spindle sources in the human cortex. Neuroscience 103, 581–592 (2001).

14. Alfonsi, V. et al. Spatiotemporal Dynamics of Sleep Spindle Sources Across NREM Sleep Cycles. Front. Neurosci. 13, (2019).

15. Vogt, B. A. & Laureys, S. Posterior cingulate, precuneal and retrosplenial cortices: cytology and components of the neural network correlates of consciousness. in Progress in Brain Research (ed. Laureys, S.) vol. 150 205–217 (Elsevier, 2005).

16. Bruner, E., Preuss, T. M., Chen, X. & Rilling, J. K. Evidence for expansion of the precuneus in human evolution. Brain Struct. Funct. 222, 1053–1060 (2017).

17. Li, R. et al. Developmental Maturation of the Precuneus as a Functional Core of the Default Mode Network. J. Cogn. Neurosci. 31, 1506–1519 (2019).

18. Kurth, S. et al. Mapping of Cortical Activity in the First Two Decades of Life: A High-Density Sleep Electroencephalogram Study. J. Neurosci. 30, 13211–13219 (2010).

19. Cajochen, C., Foy, R. & Dijk, D. J. Frontal predominance of a relative increase in sleep delta and theta EEG activity after sleep loss in humans. Sleep Res. Online SRO 2, 65–69 (1999).

20. Finelli, L. A., Borbély, A. A. & Achermann, P. Functional topography of the human nonREM sleep electroencephalogram. Eur. J. Neurosci. 13, 2282–2290 (2001).

21. Bódizs, R., Körmendi, J., Rigó, P. & Lázár, A. S. The individual adjustment method of sleep spindle analysis: Methodological improvements and roots in the fingerprint paradigm. J. Neurosci. Methods 178, 205–213 (2009).

22. Fair, D. A. et al. Functional Brain Networks Develop from a “Local to Distributed” Organization. PLOS Comput. Biol. 5, e1000381 (2009).

23. Sato, J. R. et al. Age effects on the default mode and control networks in typically developing children. J. Psychiatr. Res. 58, 89–95 (2014).

24. Sherman, L. E. et al. Development of the Default Mode and Central Executive Networks across early adolescence: A longitudinal study. Dev. Cogn. Neurosci. 10, 148–159 (2014).

25. Horovitz, S. G. et al. Decoupling of the brain’s default mode network during deep sleep. Proc. Natl. Acad. Sci. 106, 11376–11381 (2009).

26. Koike, T., Kan, S., Misaki, M. & Miyauchi, S. Connectivity pattern changes in default-mode network with deep non-REM and REM sleep. Neurosci. Res. 69, 322–330 (2011).

27. Fang, Z. et al. Sleep Spindle-dependent Functional Connectivity Correlates with Cognitive Abilities. J. Cogn. Neurosci. 32, 446–466 (2020).

28. Zhang, Z. Y., Campbell, I. G., Dhayagude, P., Espino, H. C. & Feinberg, I. Longitudinal Analysis of Sleep Spindle Maturation from Childhood through Late Adolescence. J. Neurosci. 41, 4253–4261 (2021).

29. Goldstone, A. et al. Sleep spindle characteristics in adolescents. Clin. Neurophysiol. Off. J. Int. Fed. Clin. Neurophysiol. 130, 893–902 (2019).

30. Purcell, S. M. et al. Characterizing sleep spindles in 11,630 individuals from the National Sleep Research Resource. Nat. Commun. 8, 15930 (2017).

31. Nicolas, A., Petit, D., Rompré, S. & Montplaisir, J. Sleep spindle characteristics in healthy subjects of different age groups. Clin. Neurophysiol. 112, 521–527 (2001).

32. Tarokh, L. & Carskadon, M. A. Developmental Changes in the Human Sleep EEG During Early Adolescence. Sleep 33, 801–809 (2010).

33. Tarokh, L., Van Reen, E., LeBourgeois, M., Seifer, R. & Carskadon, M. A. Sleep EEG Provides Evidence that Cortical Changes Persist into Late Adolescence. Sleep 34, 1385–1393 (2011).

34. Manoach, D. S., Pan, J. Q., Purcell, S. M. & Stickgold, R. Reduced Sleep Spindles in Schizophrenia: A Treatable Endophenotype That Links Risk Genes to Impaired Cognition? Biol. Psychiatry 80, 599–608 (2016).

35. Merikanto, I. et al. ADHD symptoms are associated with decreased activity of fast sleep spindles and poorer procedural overnight learning during adolescence. Neurobiol. Learn. Mem. 157, 106–113 (2019).

36. Plante, D. T. et al. Topographic and sex-related differences in sleep spindles in major depressive disorder: A high-density EEG investigation. J. Affect. Disord. 146, 120–125 (2013).

37. Merikanto, I. et al. Genetic risk factors for schizophrenia associate with sleep spindle activity in healthy adolescents. J. Sleep Res. 28, (2019).

38. Göder, R. et al. Impairment of sleep-related memory consolidation in schizophrenia: relevance of sleep spindles? Sleep Med. 16, 564–569 (2015).

39. Loyer Carbonneau, M., Demers, M., Bigras, M. & Guay, M.-C. Meta-Analysis of Sex Differences in ADHD Symptoms and Associated Cognitive Deficits. J. Atten. Disord. 108705472092373 (2020) doi:10.1177/1087054720923736.

40. Rock, P. L., Roiser, J. P., Riedel, W. J. & Blackwell, A. D. Cognitive impairment in depression: a systematic review and meta-analysis. Psychol. Med. 44, 2029–2040 (2014).

41. Hahn, M. et al. Developmental changes of sleep spindles and their impact on sleep-dependent memory consolidation and general cognitive abilities: A longitudinal approach. Dev. Sci. 22, e12706 (2019).

42. Reynolds, C. M., Short, M. A. & Gradisar, M. Sleep spindles and cognitive performance across adolescence: A meta-analytic review. J. Adolesc. 66, 55–70 (2018).

43. Iber, C., Ancoli-Israel, S., Chesson Jr., A.L. & Quan, S. F. The AASM Manual for Scoring of Sleep and Associated Events Rules, Terminology and Technical Specifications. (American Academy of Sleep Medicine, 2007).

44. AASM Scoring Manual - American Academy of Sleep Medicine. American Academy of Sleep Medicine – Association for Sleep Clinicians and Researchers https://aasm.org/clinical-resources/scoring-manual/.

45. Ujma, P. P., Sándor, P., Szakadát, S., Gombos, F. & Bódizs, R. Sleep spindles and intelligence in early childhood–developmental and trait-dependent aspects. Dev. Psychol. 52, 2118–2129 (2016).

46. Dustman, R. E., Shearer, D. E. & Emmerson, R. Y. Life-span changes in EEG spectral amplitude, amplitude variability and mean frequency. Clin. Neurophysiol. 110, 1399–1409 (1999).

47. Tukey, J. W. Exploratory Data Analysis. (Addison-Wesley Publishing Company, 1977).

48. Rüger, B. Das maximale signifikanzniveau des Tests: “LehneHo ab, wennk untern gegebenen tests zur ablehnung führen”. Metrika 25, 171–178 (1978).

49. Abt, K. Descriptive data analysis: a concept between confirmatory and exploratory data analysis. Methods Inf. Med. 26, 77–88 (1987).

